# Snapshots of Mitochondrial Fission Imaged by Cryo-Scanning Transmission Electron Tomography

**DOI:** 10.1101/2024.10.25.620016

**Authors:** Peter Kirchweger, Sharon Grayer Wolf, Neta Varsano, Tali Dadosh, Guenter P. Resch, Michael Elbaum

**Affiliations:** Department of Chemical and Biological Physics; Department of Chemical and Structural Biology; Department of Chemical Research Support, Weizmann Institute of Sciences, Rehovot, Israel; Nexperion e.U.-Solutions for Electron Microscopy, Vienna, Austria

## Abstract

Mitochondria undergo constant remodeling via fission, fusion, extension, and degradation. Fission, in particular, depends on the accumulation of the mitochondrial fission factor (MFF) and subsequent recruitment of the dynamin-related protein Drp1. We used cryo-scanning transmission electron tomography (cryo-STET) to investigate mitochondrial morphologies in MFF mutant cells (MFF^-/-^) in ATP-depleting conditions that normally induce fission. The capability of cryo-STET to image through the cytoplasmic volume to a depth of 1 µm provides visualization of mitochondria and their surroundings intact. We imaged changes in mitochondrial morphology and cristae structure and contacts with endoplasmic reticulum, degradative organelles, and cytoskeleton at stalled fission sites. We found disruption of the outer membrane at contact sites with endoplasmic reticulum and degradative organelles at sites of mitophagy. We identified fission sites where the inner mitochondrial membrane is already separated while the outer is still continuous. While MFF is a general fission factor, these observations demonstrate that mitochondrial fission can proceed to the final stage in its absence. The use of cryo-STET allays concerns about the loss of structures due to sample thinning required for cryo-TEM tomography.

**Summary Statement:** Imaging the whole cytosol in three dimensions greatly aids in understanding cellular processes. Here, we applied cryo-scanning transmission electron tomography to study stages of mitochondrial fission in the absence of the mitochondrial fusion factor protein.

## Introduction

Obtaining three-dimensional (3D) information on cellular organelles enables advances in cell biology, inspiring the development of various advanced imaging methods (Glancy 2020). These methods center largely on fluorescence microscopy (FM) due to the possibilities for genetic tagging with fluorescent proteins. Modern techniques bring the resolution of now-routine imaging modalities below 100 nm into the domain of intracellular organelles (e.g., Jakobs and Wurm 2014; Stephan et al. 2019; Kleele et al. 2021; Landoni et al. 2024). While fluorescence provides a powerful molecular identification in the area of interest, the unlabelled cellular theater remains dark. In contrast, classical electron microscopy (EM) methods expose the cellular architecture. Still, the harsh protocols of specimen preparation, including typically fixation, dehydration, and embedding, as well as the requirements for chemical staining, raise questions about the fidelity of the image representation. Cryo-microscopy takes an alternative approach by embedding the hydrated specimen in a vitrified state. Its application is most famous for structural biology, in which cryo-transmission electron microscopy (cryo-TEM), combined with extensive image averaging, reaches near-atomic resolution for macromolecular structures. The cryo-TEM approach was extended to tomography (cryo-ET) for studies of cells (e.g., Frank 2006; Young and Villa 2023). In parallel, other 3D cryo-microscopy techniques have emerged for studying cells, notably soft X-ray tomography (Schneider et al. 2010) and serial surface imaging using a focused ion beam (FIB) with scanning electron microscopy (SEM, Schertel et al. 2013; Hoffman et al. 2020). Each EM method has its strengths and limitations, which have been discussed elsewhere (e.g., Elbaum 2018; Varsano and Wolf 2022). Cryo-ET requires specimens of a maximum of a few hundred nanometers, significantly thinner than the typical scale of the cell and even thinner than most mitochondria. The current practice to overcome this limitation is to prepare thin lamellae by FIB milling (Marko et al. 2007; Villa et al. 2013). An alternative approach, cryo-scanning transmission electron tomography (cryo-STET), offers specific advantages (Wolf, Houben, and Elbaum 2014; Kirchenbuechler et al. 2015; Wolf et al. 2017; Kirchweger, Mullick, Swain, et al. 2023) for studies at the scale of cytoplasmic organelles. Most notable is the ability to work with much thicker specimens than conventional cryo-TEM, reaching one micron or more while retaining nanometer-scale resolution for structures not amenable to image averaging.

Several studies have used cryo-ET to study mitochondria in adherent fibroblast cells. For example, recruitment of cytoskeleton to constriction sites correlated with the depolarization of the membrane potential (Mageswaran et al. 2023). These studies show many details of the ultrastructure of mitochondria and their surroundings. However, it was possible to study only regions at the very edge of a cell or lamellae of roughly 100 nm–250 nm. Therefore, it was not feasible to characterize whole mitochondria and their surroundings intact. In this work, we apply cryo-STET to mitochondria within their intact cellular milieu to study mitochondrial fission and the effect of knock-out of the mitochondrial fission factor (MFF).

Mitochondrial homeostasis involves perpetual extension and fusion on one hand and fission and degradation on the other. Therefore, fission is essential for organelle biogenesis, quality control, metabolism, and apoptosis (Friedman and Nunnari 2014; Mattie, Krols, and McBride 2019). The core machinery involved in fission (Kraus et al. 2021) includes the GTPase dynamin-related protein 1 (DRP1) (Labrousse et al. 1999), its primary outer mitochondrial membrane (OMM) adaptor proteins MFF (Gandre-Babbe and van der Bliek 2008; Otera et al. 2010; Losón et al. 2013) or Mid49/51 (Palmer et al. 2011), together with other cellular structures such as the ER (Friedman et al. 2011; Abrisch et al. 2020) and cytoskeleton (De Vos et al. 2005; Ji et al. 2017). After recruitment to the OMM, DRP1 forms puncta at fission sites (Otera et al. 2010; Ji et al. 2015) and forms a ring around the constriction site to finalize the fission step (Kalia et al. 2018). DRP1 also binds to actin filaments, which increases the fission activity (Ji et al. 2015; Hatch et al. 2016).

MFF forms puncta on the outer mitochondrial membrane (Otera et al. 2010). Its oligomerization is specifically required for DRP1 recruitment (R. Liu and Chan 2015; A. Liu, Kage, and Higgs 2021). Observations by fluorescence microscopy reveal that deletion of MFF results in elongated mitochondria (Gandre-Babbe and van der Bliek 2008; Otera et al. 2010; Losón et al. 2013; A. Liu, Kage, and Higgs 2021) because fission is inhibited while fusion is still ongoing. In wild-type cells, the mitochondrial network can be disrupted by treatment with oligomycin (De Vos et al. 2005; Yang et al. 2018). Oligomycin binds to the proton channel in the c-ring of the ATP synthase and blocks protons from passing through, which increases the gradient (Lardy, Johnson, and McMuray 1958; Lee and O’Brien 2010; Symersky et al. 2012). Therefore, we expected that the oligomycin treatment of MFF mutant cells would reveal early intermediates in the fission process.

Using cryo-STET, we aimed to capture snapshots of early mitochondrial fission together with surrounding cellular structures that may be involved in the process. To this end, we induced fission by treating MFF^-/-^ knock-outs of mouse embryonic fibroblast (MFF^-/-^ MEFs, Losón et al. 2013) and human bone osteosarcoma cells (MFF^-/-^ U-2 OS, A. Liu, Kage, and Higgs 2021) with oligomycin. Under these fission-inducing conditions, WT cells displayed a disrupted mitochondrial network. In contrast, the mitochondrial network of MFF^-/-^ mutant cells remained elongated but appeared irregular in size and shape at the level of fluorescence microscopy. Using SRRF-enhanced cryo-FM (cryo-SRRF, Kirchweger, Mullick, Swain, et al. 2023), we located the stalled fission sites for detailed investigation by cryo-STET. Among the unusual morphologies detected, we could also observe characteristic features of normal mitochondrial fission, such as wrapping of ER around the constriction site, the presence of microtubules and actin filaments nearby, the formation of narrow OMM tubes, and OMM constriction with inner membrane separation. OMM scission, the final fission step, is a function of MFF prerequisite to DRP1 oligomerization. Still, it can clearly occur even in the absence of MFF, perhaps using MiD49/51 as adaptor proteins, or by another process such as mechanical tension by the cytoskeleton. The tomograms provide complete 3D snapshots of intact mitochondria and the surrounding cytosol, highlighting the strength of cryo-STET as a tool in cell biology.

## Results

### Mitochondrial Morphology by FM

First, we confirm the oligomycin sensitivity of the wt and MFF-/-cells using conventional FM. In wild-type MEFs (Chen et al. 2003), the reticulum of serpentine mitochondria (Fig. 1A) was fragmented by oligomycin treatment (Fig. 1B) as expected (De Vos et al. 2005; Yang et al. 2018). In MFF^-/-^ cells, elongated, less branched mitochondria were observed showing large ‘balloon-like’ structures (Fig. 1C). Under oligomycin treatment, mitochondria remain tubular as previously reported (Gandre-Babbe and van der Bliek 2008; Otera et al. 2010; Losón et al. 2013; Ji et al. 2017), but show irregularities that suggest fission sites (Fig. 1D), although the linear configuration argues against complete rupture. We aim to visualize these sites using cryo-STET at a higher resolution. Though fission does not proceed efficiently in MFF^-/-^ cells, mitochondria still show constrictions and sites apparently marked for fission, hinting that fission was initiated but not completed. Additionally, the mitochondria display various morphologies. These include extended mitochondria with a regular diameter, widened cylindrical, or beads-on-a-string morphology. In addition, several large balloon-like and fragmented mitochondria were observed (Fig. 1D).

**Fig. 1.**
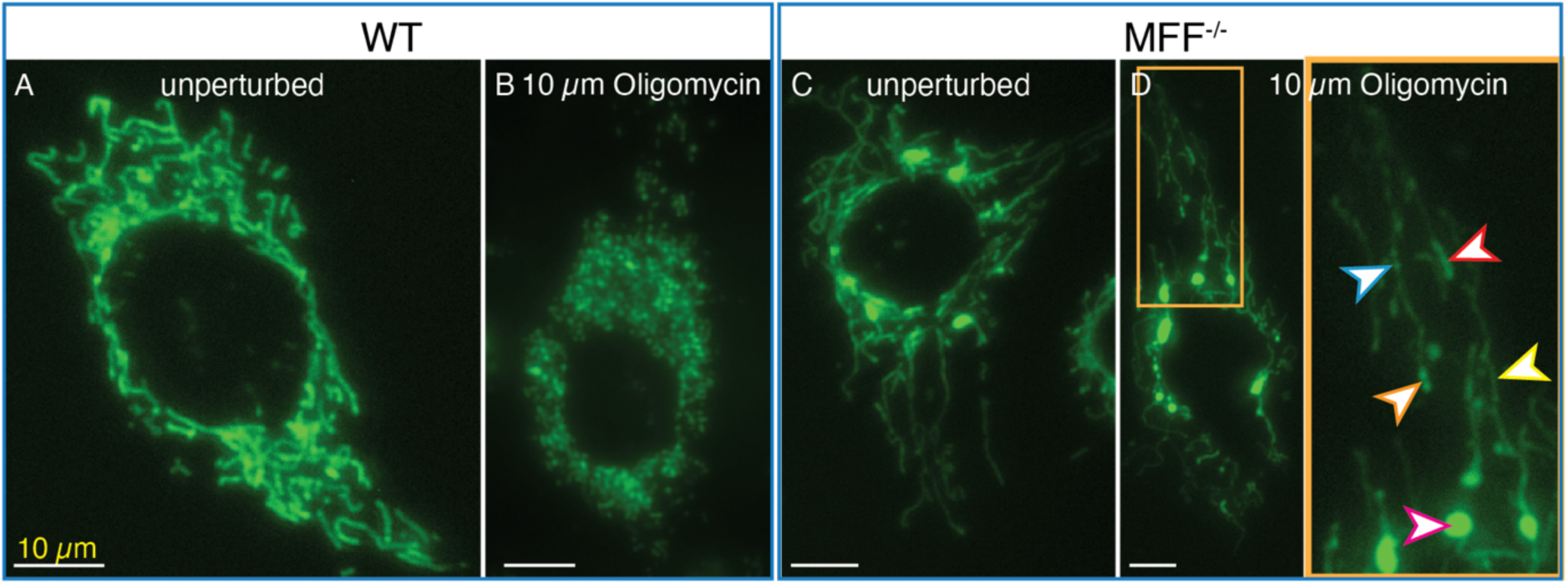
Fluorescence microscopy of (A,B) WT and (C,D) MFF^-/-^ MEFs stably expressing mitoGFP (Su9-EGFP), (A,C) unperturbed and (B,D) under fission-inducing conditions (10 µM oligomycin). Arrowheads point to different mitochondrial morphologies, namely regular (yellow), widened cylindrical (red), ‘bead-on-a-string’ (orange), mitochondrial fragments (blue), and balloon-like mitochondria (pink). The scale bars are10 µm.

### Cryo-STET of MEF MFF^-/-^ mitochondria

Next, we establish the baseline for MFF^-/-^ cells under normal growth conditions without oligomycin. Mitochondria are very long and narrow. Still, some constrictions are observed.

To visualize the effect of the MFF knockout, we imaged MFF^-/-^ cells under normal conditions using cryo-STET (Fig. 2, Fig. S1-3). To capture a variety of mitochondrial morphologies, at first, a large field of view was imaged (8 µm, Fig. 2A,B). These tubular mitochondria, with a diameter of 200 nm to 300 nm, can be observed crossing the entire field of view of 8 µm (Fig. 2A+B, Fig. S1A-C, Fig. S2). Shorter mitochondria, in the range of 2 µm – 4 µm with an increased diameter up to 2 µm, can also be observed (Fig. S1C). Therefore, elongated mitochondria are the first difference between MEF WT mitochondria imaged by others (e.g., Mageswaran et al. 2023) and the MEF MFF^-/-^ mutant. This is consistent with our fluorescence microscopy observations (Fig 1C) and those of others (Gandre-Babbe and van der Bliek 2008; Otera et al. 2010; Losón et al. 2013). Amorphous calcium-phosphate (CaP) matrix granules can be observed as dark spots (dark blue arrowheads) in the mitochondria (Wolf et al. 2017) (Fig. 2A, Fig. S1A). Interestingly, in one cell, none of the mitochondria seen in the tomograms seem to have CaPs (Fig. S1B+C), even though tomograms in Fig. 2A and Fig. S1A, and FigS1B+C are taken from the same sample. Microtubules are observed near the mitochondria (Fig. 2A+B). The cryo-FM images show the elongated nature of the MEF MFF^-/^mutant (Fig. S2A) and are overlayed onto the tomograms (Fig. S2B-D).

Next, we increased the magnification and applied dual-axis tomography to obtain further insights into the cristae network. This reveals how the cristae network is organized (Fig. 2C+D, Fig S1D, Fig. S3) and even shows individual protein densities in the cristae membrane (Fig. S3, magenta arrowheads). Interestingly, a mitochondria-derived vesicle (MDV, Sugiura et al. 2014; König and McBride 2024), with a diameter of around 130 nm, can be observed pinching off from the OMM of a swollen area of the mitochondrion; the cristae structure is degraded near the vesicle but not at a distance even in the same mitochondrion (Fig. 2C+D). Additionally, a swollen mitochondrion more than 1 µm wide can be observed, with sparse and dense cristae coexisting. Another tubular mitochondrion is seen in Fig. S1D, with a well-ordered cristae network except in the vicinity of a large vesicle. Therefore, deletion of MFF in MEF cells does not disrupt the cristae structure as a whole, but the cells show elongated mitochondria with strong internal variability.

### Cryo-STET of MFF^-/-^ cells under fission-inducing conditions

We induced fission using oligomycin to block the ATP synthase in MFF^-/-^ and recorded several cryo-STET tomograms showing elongated mitochondria with multiple constriction sites (Fig. S5) to visualize the morphological changes and to understand which cellular compartments take part in stress-induced fission. While the fission process initiates, it does not come to completion. In the following, we describe the variety of morphologies observed.

The first morphology is the pearling, or ‘bead-on-a-string’ shape (Fig. 3). The area of the cell is approximately 880 nm thick. Several mitochondria reach across the entire tomogram with a field of view of 6 µm (Fig. 3A+B). Segmentation gives an impression of the 3D nature of the data (Fig. 3B). The main bodies of the mitochondria are oblate spheroid and around 600 nm to 1 µm wide. They show degraded, rounded cristae compared to the untreated cells seen in Fig. 2C + D + Fig. S2D. Interestingly, there are no CaP deposits in these mitochondria. The beads are connected via thin tubules with a diameter of 40 nm – 55 nm. Interestingly, a bundle of microtubules runs alongside the central mitochondria (Fig 3B). Three distinct fission sites are highlighted to illustrate adjacent cellular components (Fig. 3C-E). Fission site 1 (Fig. 3C) shows ER partly wrapping around the thin fission tubule. The bundle of microtubules crosses the fission site. The fission tubule is about 650 nm long and has a diameter of around 55 nm. Fission site 2 (Fig. 3D) shows ER wrapping around the fission site over a longer distance along the fission tubule. No microtubules are seen at this site. The fission tubule is 600 nm long and 40 nm in diameter. Fission site 3 (Fig. 3E) has no notable cellular features nearby. The diameter of the fission tubule is 50 nm, and its length is around 830 nm. It seems that the IMM is already separated in this fission site.

**Fig. 3.**
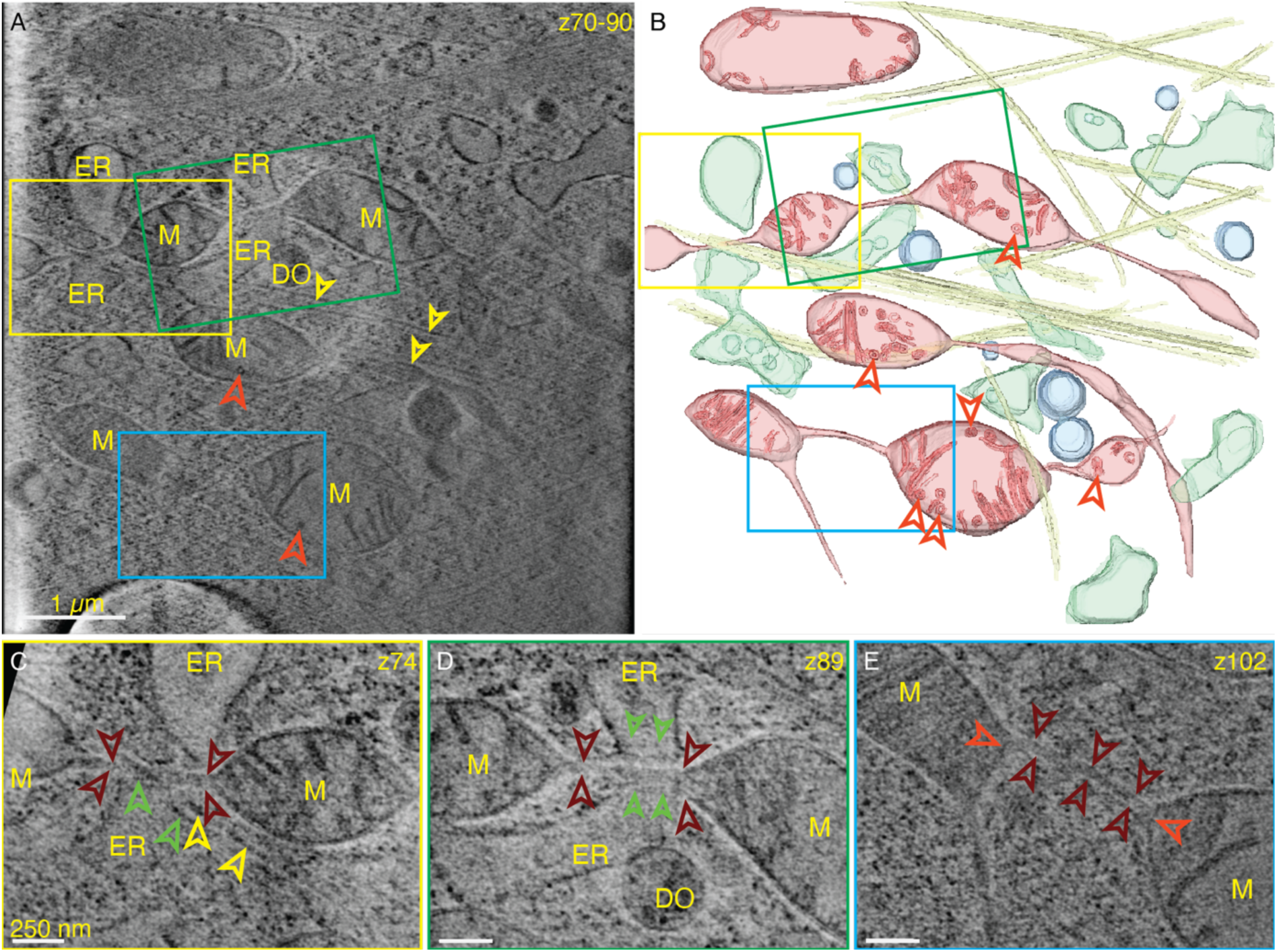
Cryo-STET tomogram of an 880 nm thick region of MFF^-/-^ MEFs under fission-inducing conditions. (A) A 60 nm thick virtual section through a tomogram showing the ‘bead-on-a-string’ topology, with its (B) 3D segmentation. Mitochondria (M, red), ER (green), digestive vesicle (DV, blue), and microtubules (yellow) are shown in the 3D segmentation. (C-E) Zoom-ins into three fission events indicated by the yellow, green, and blue boxes. Arrowheads indicate microtubules (yellow), OMM (dark red), IMM/cristae (light red), and ER membranes (green). Scale bars are 1 µm in (A) and 250 nm in (C-E).

The ‘bead-on-a-string’ morphology is also seen in U-2 OS DRP1-GFP MFF^-/-^ cells in the presence of 5 µM oligomycin. Fig. 4 shows a slice through a tomogram with several mitochondria with different morphologies near rough ER, microtubules, and lipid droplets (dr). Two mitochondrial constrictions to a diameter of about 70 nm can be observed. Zooming into one of the constriction sites (Fig. 4C) shows a microtubule running across and a lipid droplet (dr) nearby. Protein densities can be observed around the OMM. The presence of a CaP in the second constriction site suggests that transport is still active (Fig. 4D). When overlaying the cryo-FM fluorescence onto the tomogram, the DRP1-GFP signal closely resembles that of the mito-BFP (Fig. 4G). This indicates, therefore, that the DRP1 is located around the whole mitochondria rather than sitting exclusively at the constriction sites, similar to what has been reported earlier (Otera et al. 2010).

**Fig. 4.**
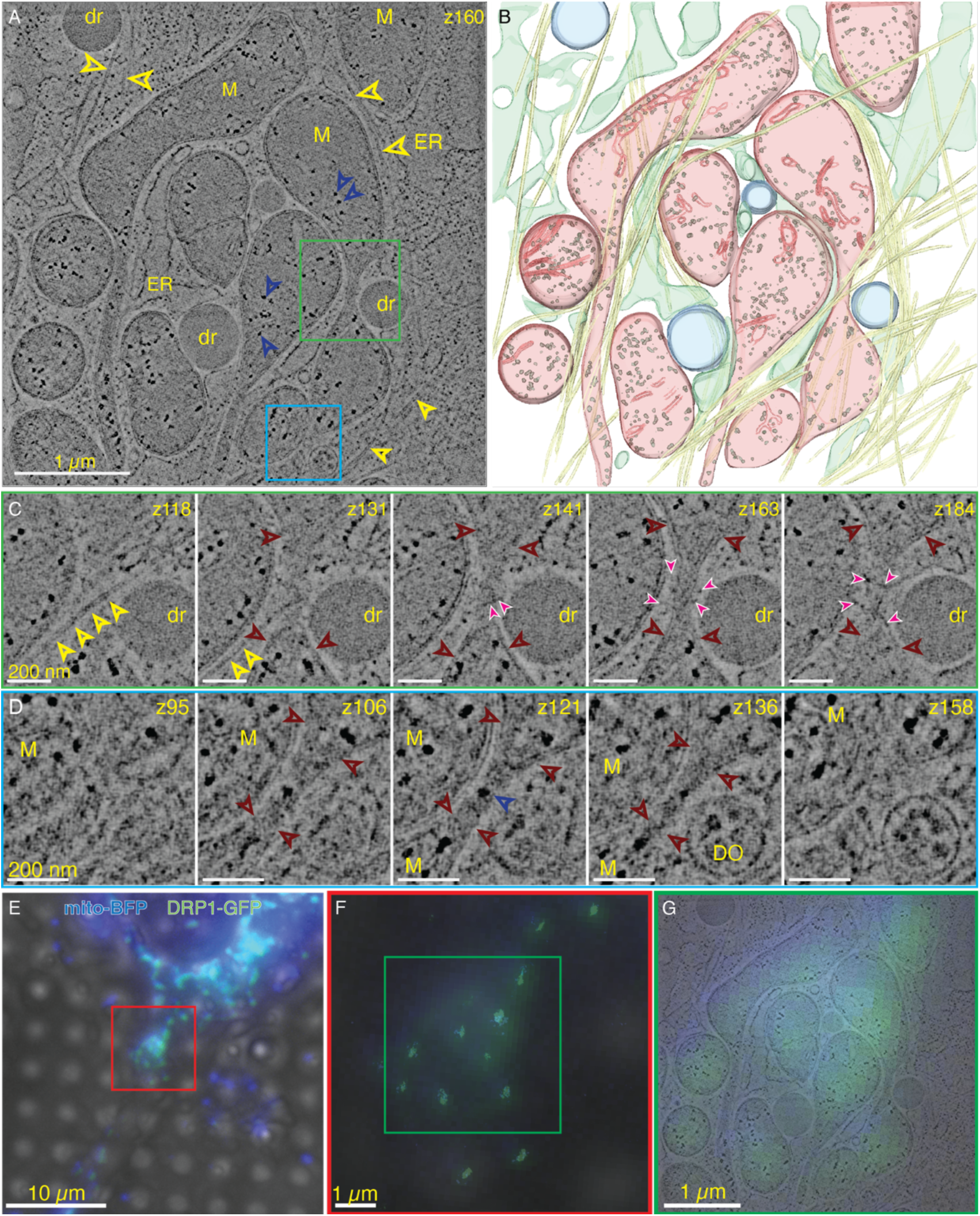
Cryo-STET tomogram of a 630 nm thick region of an MFF^-/-^ U-2 OS cell under fission-inducing conditions. (A) A single slice from the tomogram shows several mitochondria (M) with different morphologies near ER, lipid droplets (dr), and microtubules (yellow arrowheads). Dark blue arrowheads point to CaP. (B) 3D segmentation of the tomogram. Mitochondria (M, red), ER (green), lipid droplets (dr, blue), microtubules (yellow), and CaP (grey) are shown. (C) Zoom in on a constriction site indicated by the green box. The microtubules (yellow arrowheads), OMM (red arrowheads), and unknown protein densities (filled magenta arrowheads) are highlighted. (D) Zoom in on another constriction site indicated by the cyan box, highlighting a CaP (blue arrowhead) and the continuous OMM (dark red arrowheads). (E-G) Cryo-FM workflow: (E) combined maximum Z projection of widefield, mito-BFP (blue, aa1-22 of COX4 N-terminal to BFP) and Drp1-GFP (green, stable expression), (F) overlay of the cryo-SRRF (sharp peaks) onto the cryo-FM (diffuse fluorescence in the background), and (G) overlay of the cryo-FM onto the tomogram. Scale bars are (E) 10 µm, (A,F,G) 1 µm, and (C,D) 200 nm, respectively.

The second category is balloon-like mitochondria (Fig. 1D). Fig. 5 shows a cryo-STET tomogram and the respective 3D segmentation containing a 2 µm wide and 750 nm thick mitochondrion with a degraded cristae network. At the end of the mitochondrion, a fission tubule is seen, with a diameter of 40 nm extending further into the cytosol with an unclear ending. This mitochondrion does not contain any calcium phosphate granules.

**Fig. 5.**
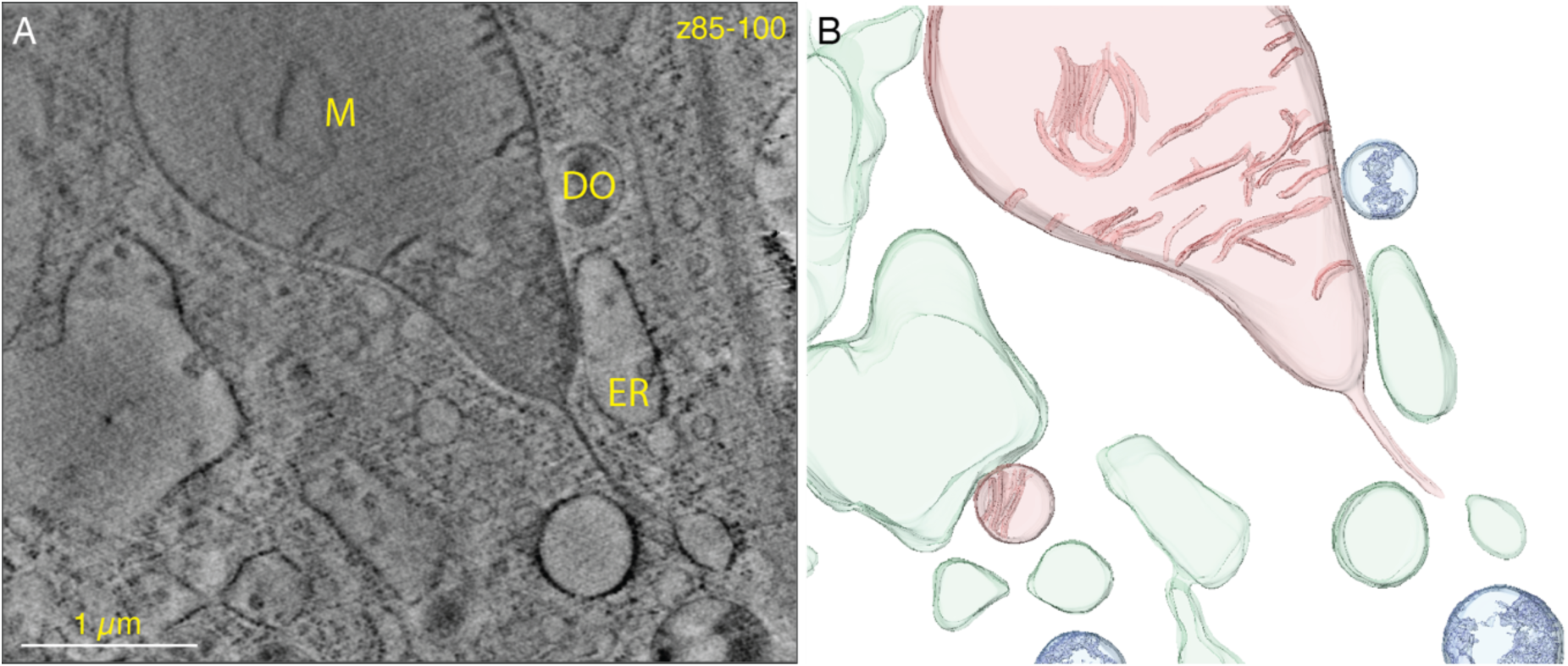
Cryo-STET tomogram of MFF^-/-^ MEFs under fission-inducing conditions in an area 730 nm thick. (A) A 48 nm thick virtual slice showing a balloon-like mitochondrion. The scale bar is 1 µm. (B) The respective 3D segmentation. Red, green, and blue in 3D segmentation show mitochondria (red), ER (green), and degradative organelles (blue).

Two other morphological features observed by FM are mitochondria with a regular and significantly increased diameter. These appear more detailed in the cryo-STET data (Fig. 6). Two neighboring dual-axis tomograms show several mitochondria, including a 10 µm long mitochondrion dividing into two (M1 + M2). The mitochondrion extends from the first tomogram into the neighboring one (Fig. 6), as seen by the overlay of the tomograms onto the cryo-FM data (Fig. S5A). This mitochondrion can be separated into three parts, displaying two stages of mitochondrial fission (Fig. S5B). In the ‘first part’ (Fig. S5B), an increased diameter of around 750 nm with a degraded cristae network is observed at one end (Fig. 6A). Roughly in the center of the first tomogram, ER is seen to wrap around the mitochondrion (Fig. 6A). Interestingly, ER is present at the top and bottom of the mitochondrion (Fig S5B, S6B). This marks the beginning of the ‘second part’ (Fig. S5B, green box), where the diameter is reduced. Interestingly, hardly any cristae can be observed in this narrower part of the mitochondria. Together, the present ER, the reduction in diameter, and the change in cristae appearance could suggest a very early stage of fission. The second part stretches further into the neighboring tomogram (Fig. 6) until a fission site in the final stage is observed, which marks the beginning of the ‘third part’ (Fig. S5B). M2 has a diameter of 250 µm – 400 µm and shows well-ordered cristae (Fig. 6C, D). A longer mitochondrion (M3) with degraded cristae also reaches into the first tomogram (Fig. 6A+B). Smaller mitochondrial fragments are observed (M4-M9), some entirely depleted of cristae (M4, M9).

**Fig. 6.**
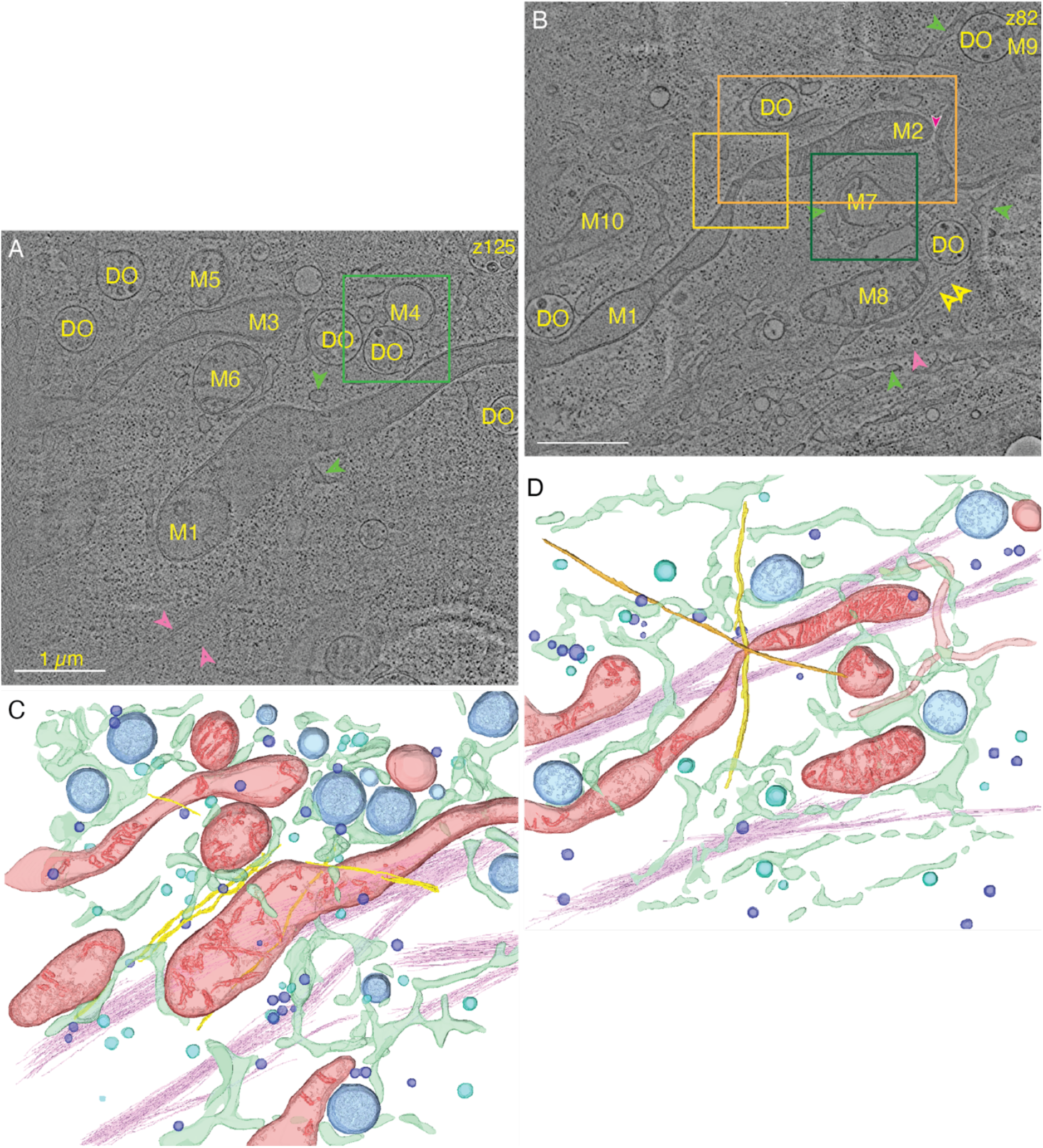
Cryo-STET tomograms of two regions of MFF^-/-^ MEFs under fission-inducing conditions, with a thickness of 750 nm and 550 nm, respectively. (A+B) A slice of two neighboring dual-axis tomograms showing several mitochondria (numbered M1-10) with their interactions with organelles and cytoskeleton. Arrowheads show ER (green), actin bundles (pink), microtubules (yellow), and protein tether between the unknown tubular structure and mitochondria (filled magenta). The overlay onto the cryo-EM image and the spatial arrangement are shown in Fig. S5. Yellow and green boxes show zoom-ins highlighted in Fig. 7 and Fig. 8. Scale bars are 1 µm. (C+D) 3D-segmentation of the tomograms. Mitochondria (red), ER (green), degradative organelles (blue), exosomes/microvesicles (purple), actin filaments (pink), unknown tubular structure (light red), other vesicles (cyan) and microtubules (yellow+orange).

### Contacts with other cellular components

After describing the changes in morphology, we highlight some contacts between mitochondria and other cellular components observed in these tomograms. In the following, we highlight some examples, even though more can be found when scrolling through the 3D volumes. Two microtubules are seen to interact with the mitochondrion at the fission site, separating M1 from M2 (Fig. 7A) and crossing on the top and bottom sides of the fission site. This fission site is particularly interesting because the IMM has already separated at two positions (Fig. 7A), although the outer membrane is still shared. This suggests that it is a fission site in a late stage. Additionally, actin bundles run alongside the mitochondrion from one tomogram into the second (Fig. 6, pink bundles). At the third part (M2, Fig 7B), whose morphology is closest to normal, these actin bundles directly interact with the mitochondrion before and after the fission site and at the very tip of M2 (Fig. S7).

**Fig. 7.**
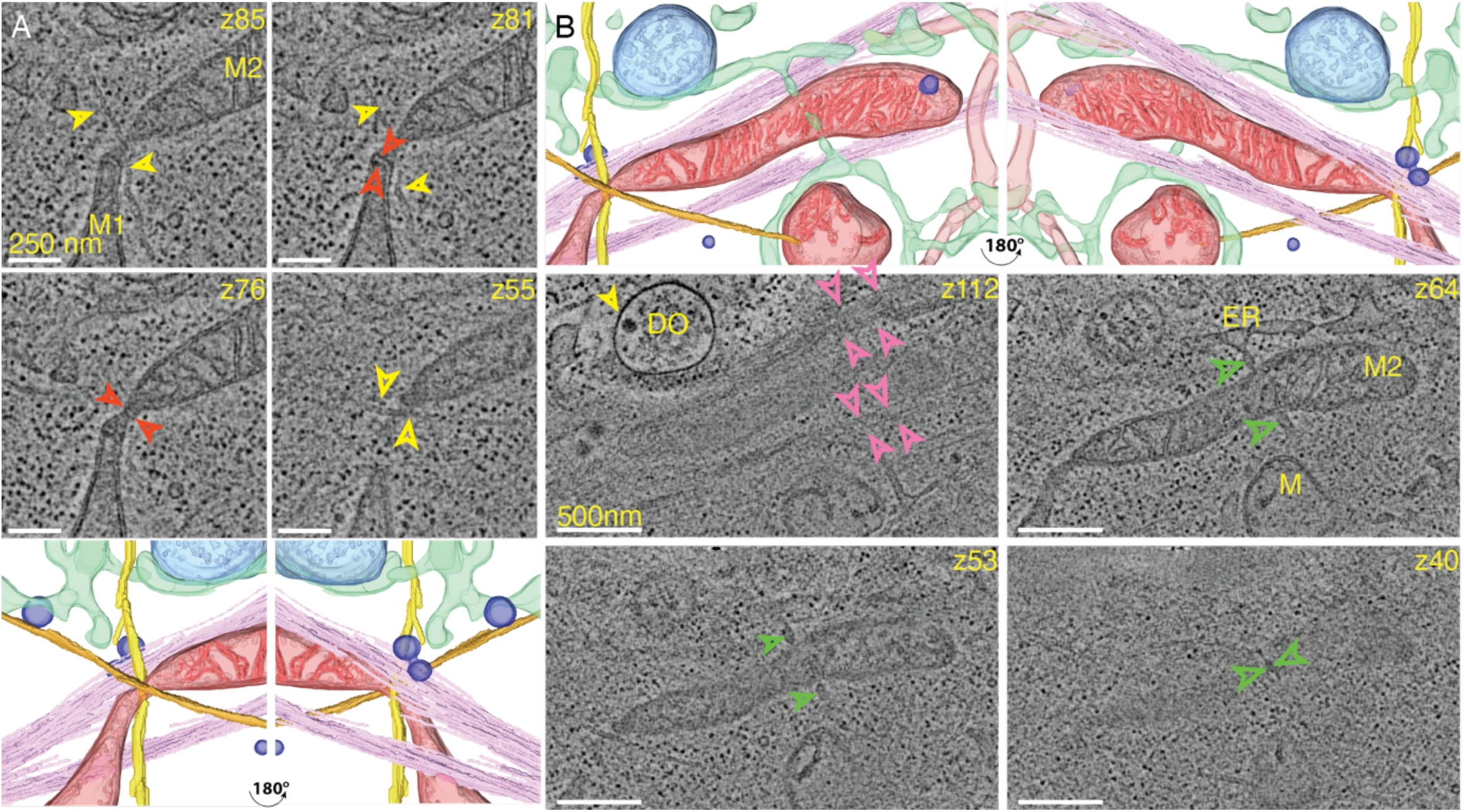
MFF^-/-^ MEFs under fission-inducing conditions. Focus on the (A) fission site and the (B) more ordered cristae network highlighted in Fig. 6 (yellow boxes). (A) A fission site where the mitochondria are separated. The IMM is already separated at two positions (red arrowheads), and microtubules are running across the fission site on the top and bottom of the mitochondrion (yellow arrowheads and yellow and orange in the segmentation). (B) 3D segmentation and slices through the tomogram showing a close-up of the actin bundles (pink arrowheads) and the ordered cristae network. ER (green network and arrowheads) is seen to reach over the mitochondria. Scale bars are 250 nm in (A) and 500 nm in (B). 3D segmentation shows mitochondria (M, red), ER (green), degradative organelles (blue), actin filaments (pink), unknown tubular structures (light red), and microtubules (yellow and orange). Red, green, yellow, and pink arrowheads point to IMM, ER, microtubules, and actin filaments, respectively.

ER is wrapped around the long mitochondrion at several positions in these two tomograms (Fig. 6, Fig. S5+Fig. S6). One such contact site appears at the tip of M1 (Fig. 6A, Fig. S6A). Two additional ER tubules are seen on top of the mitochondria (Fig. S6B, z39+48), one of which wraps around the contact (Fig. S6B, z156) in the aforementioned early-stage fission site. Another ER tubule reaches below the same mitochondrion in the ‘second part’ (Fig. S6C), and yet another appears above the center of the ‘third part’ (Fig. S6D).

A cradle-like membrane contact coincides with disrupted OMM and disrupted cristae (Fig. 8A). Some protein densities making contact between the ER and the M7 are highlighted. An unknown filled tubular structure (light red tube in Fig. 6D) that makes contact with the end of M2 (Fig. 6C+D) appears close to M7 and makes contact with the ER (Fig. 8A). This tube has a diameter between 70 nm – 80 nm. Atypically for ER, it is filled with some density. We can suggest it may be a tubulated form of a degrative organelle (Bohnert and Johnson 2022). Round degradative organelles resembling multivesicular bodies or endosomes (Fig. 8B, Fig. S6E) appear adjacent to mitochondria that lack cristae (M4, M9) and show ruptured OMM. Some densities are observed in the area where the OMM disintegrates. Interestingly, this small mitochondrial fragment retained its green fluorescence (Fig. S5), suggesting that the mitochondrial matrix is not yet acidified due to its contact with the DO (Sargsyan et al. 2015).

**Fig. 8.**
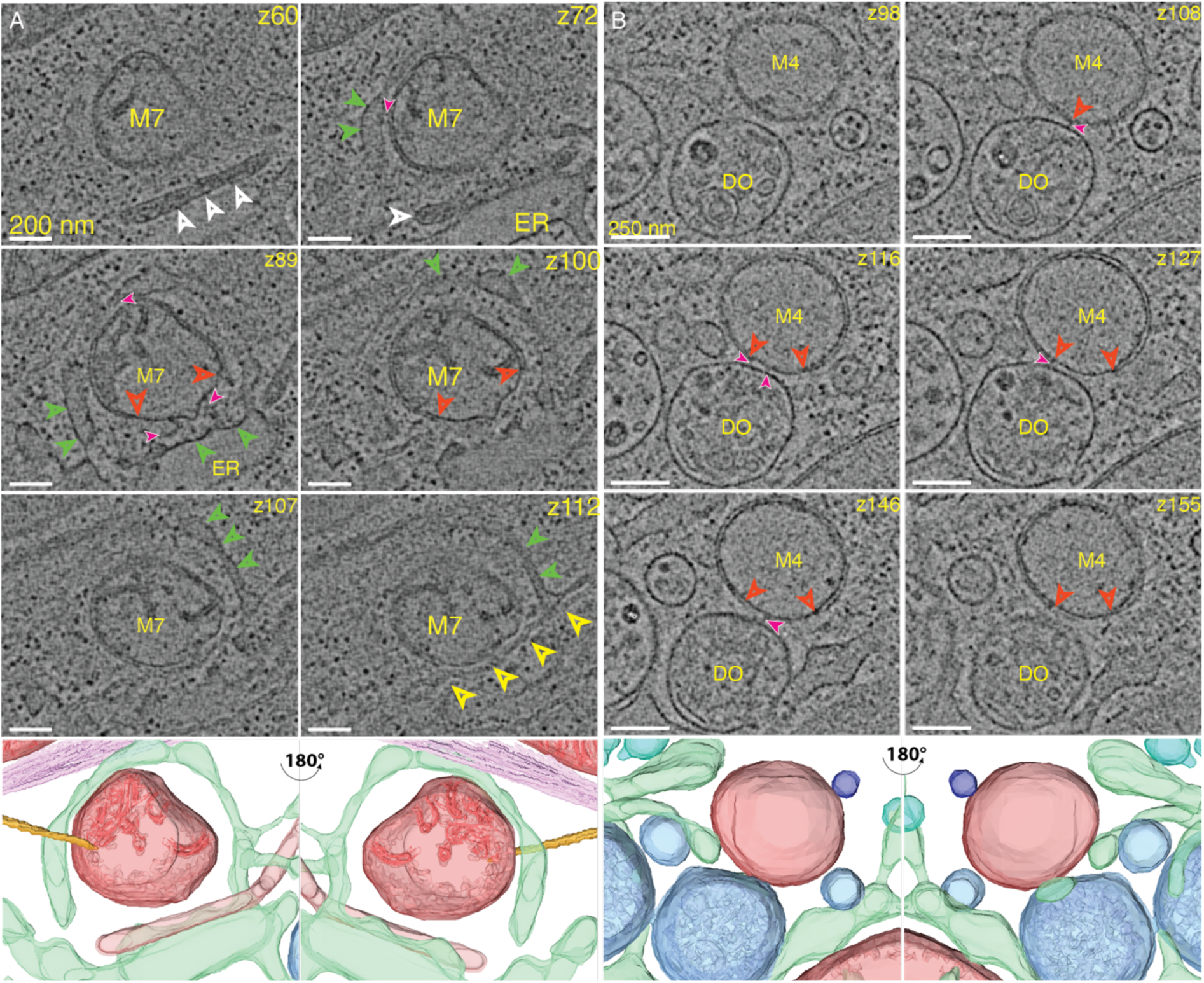
Fragmented mitochondria contact (A) ER membranes and (B) a digestive vesicle highlighted in Fig. 6 (green boxes). (A) A fragment of a mitochondrion encircled by an ER tube and the unknown tubular structure is seen in the proximity of the mitochondria (white arrowheads, z60, z72, + light red tube in the 3D segmentation). Scale bars are 200nm. (B) A mitochondrial fragment in contact with a digestive vesicle. Some observed tethering protein densities are highlighted (filled magenta arrowheads). Scale bars are 250 nm. 3D segmentation shows mitochondria (M, red), ER (green), degradative organelles (blue), unknown tubular structure (light red), and microtubules (orange). Arrowheads point to the IMM close to disrupted OMM (red), ER (green), microtubules (yellow), and unknown protein densities (filled magenta).

## Discussion

Mitochondrial morphology is a classic topic in cell biology (Lewis and Lewis 1915; Monzel, Enríquez, and Picard 2023; Preminger and Schuldiner 2024). Mitochondria undergo constant fission and fusion, dependent on protein factors for membrane remodeling (Pagliuso, Cossart, and Stavru 2018; Kraus et al. 2021). In the case of fission, MFF was identified as an early-stage adapter to the scission protein Drp1 (Otera et al. 2010). MFF is thought to oligomerize on the mitochondrial outer membrane, reducing its diameter to roughly 100 nm (R. Liu and Chan 2015; A. Liu, Kage, and Higgs 2021), at this point, Drp1 can form rings that ultimately cleave the membrane constriction (Kalia et al. 2018). Other factors, such as the Mid49/51 (Palmer et al. 2011), INF2 (Hatch, Gurel, and Higgs 2014), ER (Friedman et al. 2011), membrane tension due to cytoskeleton pulling (Mahecic et al. 2021), or other mechanical forces (Helle et al. 2017; Romani et al. 2022), have also been implicated in constricting the future cleavage site (Kraus et al. 2021). Using correlative-light and electron microscopy (CLEM, Schwartz et al. 2007)-guided cryo-STET imaging, we found that OMM constriction can progress to a very late stage even in an MFF-knockout environment. The CLEM approach, as we demonstrate here, can provide this via specific molecular fluorescence (Fig. 4E-G) or morphologies at a larger scale (Fig. 2, S5A).

Fluorescence microscopy showed that MFF^-/-^ MEF cells possess an extended mitochondrial network. Unlike wild-type, several swelled, balloon-like extensions are seen in the perinuclear region, apparently at the ends of the tubular mitochondria. Unstressed mitochondria in MFF^-/-^ cells are highly elongated and appear somewhat narrower than normal (Fig. 1C, 2, S1, S2). The cristae pattern seems well-ordered along much of the mitochondria length but is degraded in swollen areas (Fig. S1D, Fig. S3). While shorter and swollen mitochondria are observed (Fig. S1C), constrictions of mitochondria are difficult to find. Upon treatment of the MFF^-/-^ cells with oligomycin, the frequency and morphology of the constrictions change drastically. We then targeted these constrictions by cryo-FM/cryo-SRRF and cryo-STET to visualize the stalled fission sites and to elucidate the contributions of other intracellular structures at different stages.

Oligomycin treatment transformed the narrow mitochondrial cylinders of MFF^-/-^ cells into a varicose, beads-on-a-string morphology of bulges interspersed by very narrow tubules. The cristae network is also less regular in the bulges than that seen in the tubes of untreated cells, showing degraded, circular cristae (Fig 3A,B). Very thin, narrow fission tubules connect the bulges (Fig 3C-E). These appear, at least in some cases (Fig 3E), to be composed only of OMM, leaving a closed IMM on either side. This is in contrast to normal mitochondrial fission (Mageswaran et al. 2023), where the two membranes pinch together simultaneously. We can suggest that the fission tubule in the MFF^-/-^ cells may be drawn out from the isolated pinch point by a pulling force due to a bundle of microtubules running alongside the mitochondria (Fig 3A,B) and passing across one of the fission sites (Fig 3C). Rupture of the fission tubule leaves an isolated mitochondrion, for example, seen in Fig 5, where the tubule remains attached on one side.

Two other fission sites are reported here, one in the very early stage, where a change in morphology, colocalization with ER, and a crista perpendicular to the OMM was observed (Fig 6A, S5, between parts one and two), and another one at a late stage, where the IMM had already clearly separated while the OMM remained continuous (Fig 6B,7A). Additionally, in MFF^-/-^ DRP1-GFP U2-OS cells, DRP1 seems to be distributed across the whole mitochondrion and not localized at the fission sites (Fig 4E-G), consistent with an earlier report (A. Liu, Kage, and Higgs 2021).

Whereas MFF was also considered an early-stage factor in mitochondrial fission by getting recruited to ER-mitochondria contact sites (Friedman et al. 2011; Abrisch et al. 2020), we observe that fission progresses to late stages even in its absence. Early functions may be fulfilled by redundant factors, in particular Mid49/51. Even without MFF, fission initiates with recruitment of ER and microtubules to the fission sites. The fission process continues with IMM being separated already, but the cutting of the OMM by DRP1 does appear to depend on MFF in an obligatory way. This would explain the transformation of the pinch site into a long fission tubule. The final rupture could then be affected by a mechanical force, where via an anchoring factor, one end of the mitochondria gets fixed to the cytoskeleton, e.g., microtubules (Fig 3) or actin (Fig 6, 7B), while the other end gets pulled away. This bears a similarity to the formation of mitochondrial nanotunnels (Wang et al. 2015; Vincent et al. 2017)

Technologically, the cryo-FM/cryo-STET workflow combines the specificity of FM with the possibility of imaging the whole cellular environment to a thickness of around 1 µm in a near-native state without the need for sample thinning and staining. Locating the rare constriction sites without guidance from the cryo-FM would be challenging. Moreover, the fission tubules are so narrow that the probability of finding them in a FIB-milled section of 100 nm is slim. Tubules that end on one side, such as the one seen in Fig 5, might also be lost in sectioning, so such features would have to be observed multiple times for confidence in interpretation. Here, the requirement for a large statistical sampling is replaced by the ability to image through the entire cytoplasm. The lower resolution of cryo-STET, on the other hand, makes it impractical for sub-tomogram averaging so far, at least with the simple bright-field modality employed here. In recent years, we have enhanced the STEM technology (Wolf, Houben, and Elbaum 2014; Kirchweger, Mullick, Wolf, et al. 2023) by adding 3D deconvolution (Waugh et al. 2020) combined with dual-axis tomography (Kirchweger, Mullick, Swain, et al. 2023) to the toolbox. Further developments are underway, especially with 4D STEM methods (Seifer et al. 2024), and we can look forward to enhanced resolution in the foreseeable future.

## Materials and Methods

### Cell growth

MEF-WT (Chen et al. 2003) and MEF-MFF^-/-^ cells (Losón et al. 2013) expressing an eGFP in the matrix of the mitochondria with a SU9 mitochondrial targeting sequence were a gift from the Chan lab (California Institute of Technology, USA). U-2 OS-WT and U-2 OS DRP1GFP-MFF^-/-^ (Ji et al. 2017; A. Liu, Kage, and Higgs 2021) cells were gifts from the Higgs Lab (Dartmouth, Hanover, NH, USA). MEF and U-2 OS cells were maintained in DMEM (Gibco, 41965) + 10 % FCS + 4 mM L-Glu + 1 % Pen/Strep + 1 mM Sodium Pyruvate (Biological Industries, 03-042-1B) at 37 ℃ and 5 % CO_2_ in a humid environment. The U-2 OS cells were transfected with 800 ng of mitoBFP (a gift from the Higgs Lab) in 1 mL volume in a 3.5 cm dish (Ji, Hatch, et al. 2015) using jetPRIME (Polyplus) according to the short DNA transfection protocol. To induce fission, the cells were incubated with 5 µM – 10 µM oligomycin (Sigma: O4876) for around 3 h before imaging or vitrification.

### Fluorescence Microscopy

For FM imaging, the MEF cells were grown on a 3.5 cm cell culture dish and treated as described above. The images were recorded on an Olympus IX51 microscope equipped with an Olympus U-LH100L-3 camera.

### Specimen preparation

For specimen preparation, the cells were grown on either R3.5/1 200 mesh grids (Quantifoil), onto which 10nm of continuous carbon was floated, or G200F1 R2/2 SiO2 grids (Quantifoil). The grids were glow discharged and coated with human fibronectin (FAL356008, Corning, 50 µg µL−^1^) in PBS without Mg^2+^/Ca^2+^ for about 1 h. Subsequently, 2 mL of growth medium was added, and the cells were seeded on the grids. The cells were grown in the incubator to a density of about 1-2 cells per grid square, usually overnight. Oligomycin was added roughly three hours before chemical fixation or plunging. Immediately before plunge freezing, MEF cells in Fig 6 were chemically fixed using 4 % paraformaldehyde and 0.1 % glutaraldehyde in PBS buffer. The U-2 OS cells were transfected and plunge-frozen after around 18 h of transfection. The U-2 OS cells were not chemically fixed. Vitrification was performed as described (Kirchweger, Mullick, Swain, et al. 2023). In brief, we used a Leica EM GP plunger (Leica Microsystems, Vienna, Austria, Resch et al. 2011). The chamber was set to 37 ℃ and 95 % humidity. Before blotting, 3 µL of growth medium (front side) and 1.5µL of homemade 15nm gold fiducial markers (back side) were added. Subsequently, the grids were blotted from the back side for 5 sec and plunged into liquid ethane.

### Cryo-FM data recording and analysis

Cryo-FM grid maps were recorded on a Leica Cryo CLEM as described using the GFP channel (Leica Microsystems, Schorb et al. 2017; Kirchweger, Mullick, Swain, et al. 2023). The microscope is equipped with a Hamamatsu Orca-Flash 4.0 sCMOS camera. A focus map was recorded in the GFP channel, and the subsequent grid map was recorded as a Z-stack of typically 20µm in the BF and GFP channels. Cryo-SRRF data was acquired as described (Kirchweger, Mullick, Swain, et al. 2023). In short, an area of interest was chosen, and 500 frames of around 1024×1024 pixels of different z-heights were recorded. The data was analyzed using the NANOJ-SRRF (Gustafsson, Falkenberg, and Larsson 2016; Culley et al. 2018) plugin in FIJI (Schindelin et al. 2012) using a Python script to find the best SRRF settings by splitting the stack into even and odds and calculating the FRC of the final SRRF images. The script is available on Github: (https://github.com/PKirchweger/SRRF-macro).

### Cryo-STET data collection

We followed the previously reported protocol for the alignment and set-up of the microscope (Kirchweger, Mullick, Wolf, et al. 2023; Kirchweger, Mullick, Swain, et al. 2023). Bright Field (BF)-cryo-STET data was collected on a Titan Krios G3 (Thermo Fisher Scientific), operated at 300 kV, equipped with an X-FEG electron source, a dual-axis stage, and a Fischione high-angle annular dark-field (HAADF) detector. We used a semi-convergence angle of 1.2 mrad, a 70 µm condenser aperture, and spot size 9, which resulted in a current of roughly 20 pA of the factory-calibrated projection screen. BF-cryo-STET collection on the HAADF was achieved by inserting a 20 µm objective aperture to limit the collection angles to 3mrad and, using diffraction alignment, moving the beam off-axis onto the sensitive area of the HAADF (Kirchweger, Mullick, Swain, et al. 2023). Cryo-STET data were collected using the STEM Tomography software (Fig. 2A, Fig. 3, Fig. 5; Thermo Fisher) or SerialEM (Fig. 2C, Fig. 4, Fig. 6) (Mastronarde 2003, 2005; Resch 2019).

In the case of the STEM Tomography software, a low-resolution Atlas was recorded, and positions for tilt-series data collection were chosen using the fluorescent map as a guide. Tilt series were recorded in a bidirectional manner, from −20° to 60°, and then from −22° to −60°, with an image of 2048×2048 pixels at a magnification of 21000× or 29000×, with sampling steps of 4.212nm or 3.001nm, respectively. The holder calibrations were used to skip autofocusing during the data collection.

In the case of SerialEM, cryo-STET data collection is described previously (Kirchweger, Mullick, Swain, et al. 2023; Kirchweger, Mullick, Wolf, et al. 2023). Briefly, a low-resolution map was recorded in LM-STEM mode (260× magnification), to which the fluorescent map was registered. Medium-resolution square maps at 3500× magnification were recorded based on how the cell is positioned on the grid square and the fluorescence of the mitochondria. Then, low-dose mode was activated, and tilt series positions were defined using Anchor Maps and recorded in View and Preview settings. Tilt series were recorded starting at 0°, using the dose symmetric tilt scheme up to +/-60^◦^ (Hagen, Wan, and Briggs 2017). Tilt series with 4096×4096 pixels were recorded at 29000× or 41000× magnification, resulting in sampling steps of 1.447nm and 1.021nm, respectively.

A SerialEM script was developed for dual-axis cryo-STET data collection (see supplementary information). The script involves several steps. First, a low-resolution Record image of the center of the grid is recorded and saved as a map in the navigator. Then, the grid is rotated by 90°, and a second image at the center is recorded and saved as a navigator map. Then, the Align-with-rotation procedure is called, and the two center images are compared and aligned. The resulting rotation and shift are applied to all navigator items, including the Map images. After terminating the script, the tilt-series data collection is set up for the second axis. Realign-to-item should work well with the rotated Anchor Maps if the tilt series position is not too close to the square’s edge.

### Tomogram reconstruction and combination

Tilt series alignment was performed in IMOD (Kremer, Mastronarde, and McIntosh 1996; Mastronarde and Held 2017). Patch tracking was used for the tomogram in Fig. 3, where no fiducials were present. Other tilt series were aligned using fiducials. All tomograms were reconstructed to an image size of 2048×2048 pixels. For visualization and dual-axis tomogram combination, tomograms were reconstructed using the SIRT-like filter with constant=30, while the weighted back projection with a HammingLikeFilter set to 0.0 was used for subsequent deconvolution (ERDC, Kirchweger, Mullick, Swain, et al. 2023).

### ERDC

For ERDC, gold beads were removed using the findbeads3D pipeline in IMOD. ERDC was performed using the recently published Python script as described (Waugh et al. 2020; Croxford et al. 2021; Kirchweger, Mullick, Swain, et al. 2023). The script is available on GitHub (https://github.com/PKirchweger/CSTET_Deconvolution).

### Segmentation

Amira 3D software v2022.2 was used for segmentation and 3D representation of the reconstructed data (Thermo Fisher Scientific, Waltham, MA, USA). Distance analysis was performed using Amira 3D’s ’Surface Distance’ module, which measures the distance between two triangulated surfaces. The module calculates the nearest point on the opposite surface for each vertex and outputs vector magnitudes. These distances are visualized with color intensity representing the distance at each node.

## Supporting information

Supplemental Images and Text

## Acknowledgments

MEF-“WT,” MFF^-/-^ cells were a gift from the Chan Lab, California Institute of Technology. The U-2 OS cells were a gift from Higgs Lab, Geisel School of Medicine, Dartmouth. The authors thank Tal Ilani, Department of Chemical and Structural Biology, Weizmann Institute of Sciences, for her support during the studies.

## Conflicts of interest

“The authors declare no conflict of interest.”

## Author contributions

Conceptualization, P.K., S.G.W., and D.F.; methodology, P.K., D.F; software, G.R., S.G.W, P.K; validation, P.K., S.G.W., and M.E.; investigation, P.K., SG.W.; resources, P.K., S.G.W, D.F; data curation, P.K.; writing—original draft preparation, P.K.; writing—review and editing, all authors; visualization, P.K., N.V, M.E.; supervision, D.F. and M.E.; project administration, S.G.W., D.F. and M.E.; funding acquisition, P.K., S.G.W, D.F and M.E.

## Funding

P.K. was funded by the Austrian Science Fund (FWF) through a Schrödinger Fellowship J4449-B. For the purpose of open access, the authors have applied a CC-BY public copyright license to any Author Accepted Manuscript version arising from this submission. M.E. and S.G.W. acknowledge funding from the Israel Science Foundation, grant no.1696/18, and the European Union Horizon 2020 Twinning project, IMpaCT (grant no.857203). Funding from the ERC project CryoSTEM (grant no. 101055413) is also acknowledged. Views and opinions expressed are, however, those of the author(s) only and do not necessarily reflect those of the European Union or the European Research Council Executive Agency. Neither the European Union nor the granting authority can be held responsible. M.E. is the incumbent of the Sam and Ayala Zacks Professorial Chair and head of the Irving and Cherna Moskowitz Center for Nano and Bio-Nano Imaging. The laboratory of M.E. has benefited from the historical generosity of the Harold Perlman family.

## Data availability

The tomograms presented here are available on the EMDB:

The cryo-SRRF and the deconvolution Python script are available on GitHub under https://github.com/PKirchweger/SRRF-macro and https://github.com/PKirchweger/CSTET_Deconvolution.

The dual-axis SerialEM script is available at https://serialemscripts.nexperion.net/script/81.

**Table 1.**
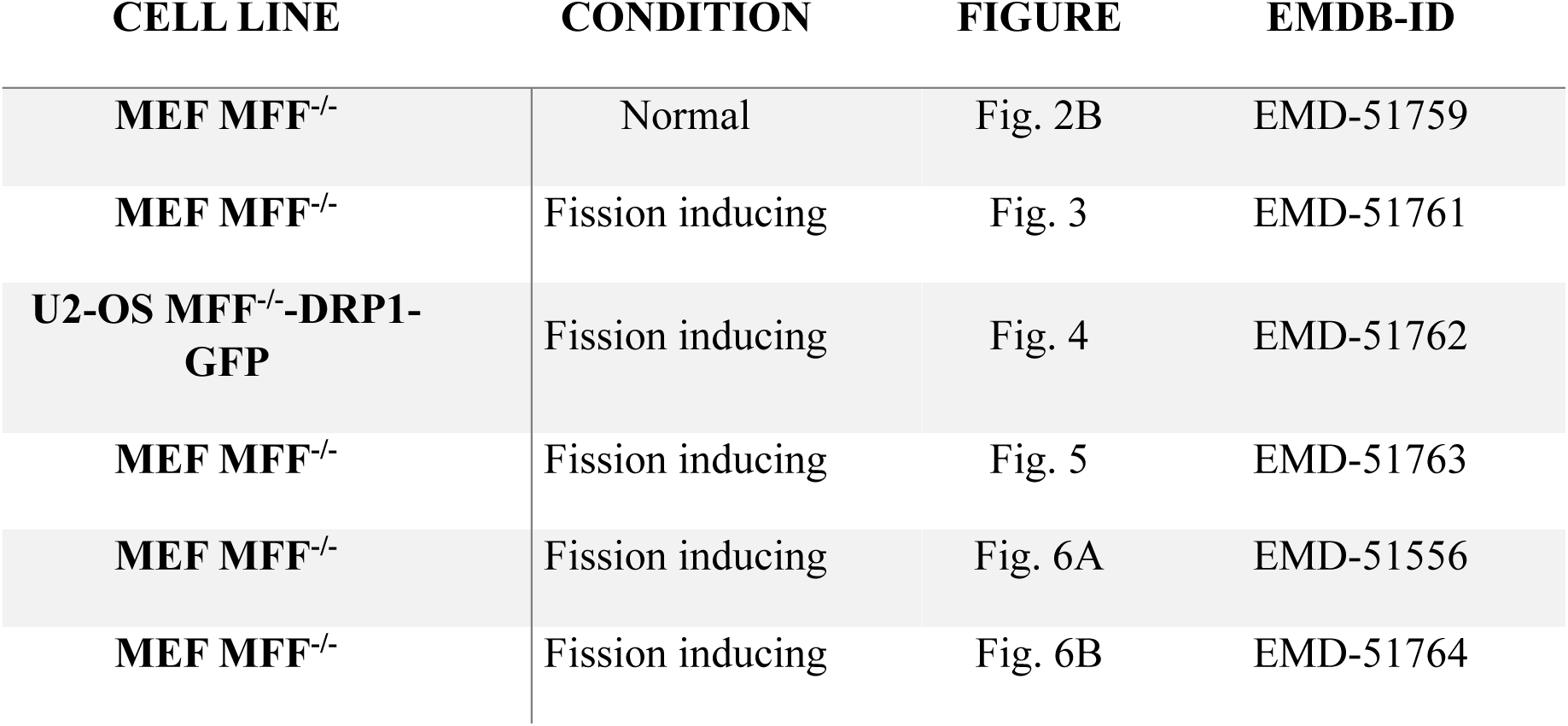
Tomograms uploaded to the EMDB.

## Notes

### Competing Interest Statement

The authors have declared no competing interest.

