## Supplemental Images and Text for "Snapshots of Mitochondrial Fission Imaged by Cryo-Scanning Transmission Electron Tomography"

### Supplemental information
# -
### Stalled Mitochondrial Fission Imaged by Cryo-Scanning Transmission Electron Tomography

Peter Kirchweger[1,2,*], Sharon Grayer Wolf[3], Neta Varsano[3], Tali Dadosh[3], Guenter P. Resch[4], Michael Elbaum[1]

1 Department of Chemical and Biological Physics, 2 Department of Chemical and Structural Biology, 3 Department of Chemical Research Support, Weizmann Institute of Sciences, Rehovot, Israel, 4 Nexperion e.U.-Solutions for Electron Microscopy, Vienna, Austria

### 1 Supplemental Figures

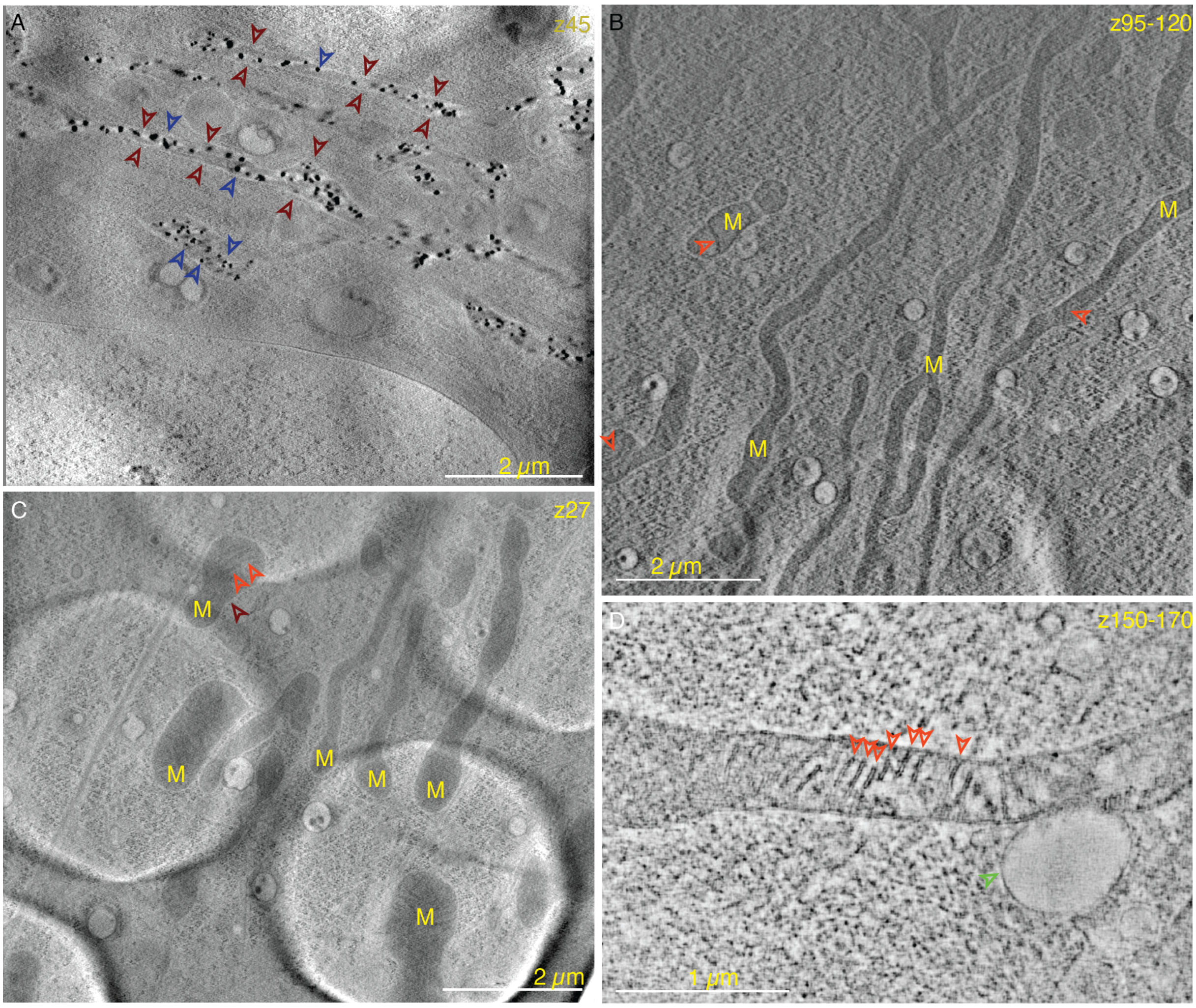


**Fig. S1.** Gallery of MFF$^{-/-}$ Mitochondria without oligomycin. (A-C) Single virtual slices through lower-resolution tomograms of mitochondria (M) from two different cells on the same specimen. Mitochondrial morphologies include aggregation (A), elongated mitochondria (B), and shorter mitochondrial fragments (C). Black dots in (A) show the CaPs (blue arrowheads). Cristae in some mitochondria, even the cristae, are visible (light red arrowheads). The OMM (dark red arrowheads) highlight the OMM. The circular structures seen very clearly in (C), with a diameter of around 3.5 nm, are holes in the carbon support layer. Scale bar is 2 µm. (D) A 40 nm thick virtual slice through a higher-resolution dual-axis tomogram. The mitochondrion interacts with a big vesicle (light green arrowhead). The cristae structure (light red arrowheads) is very clearly visible. Scale bar is 1 µm.

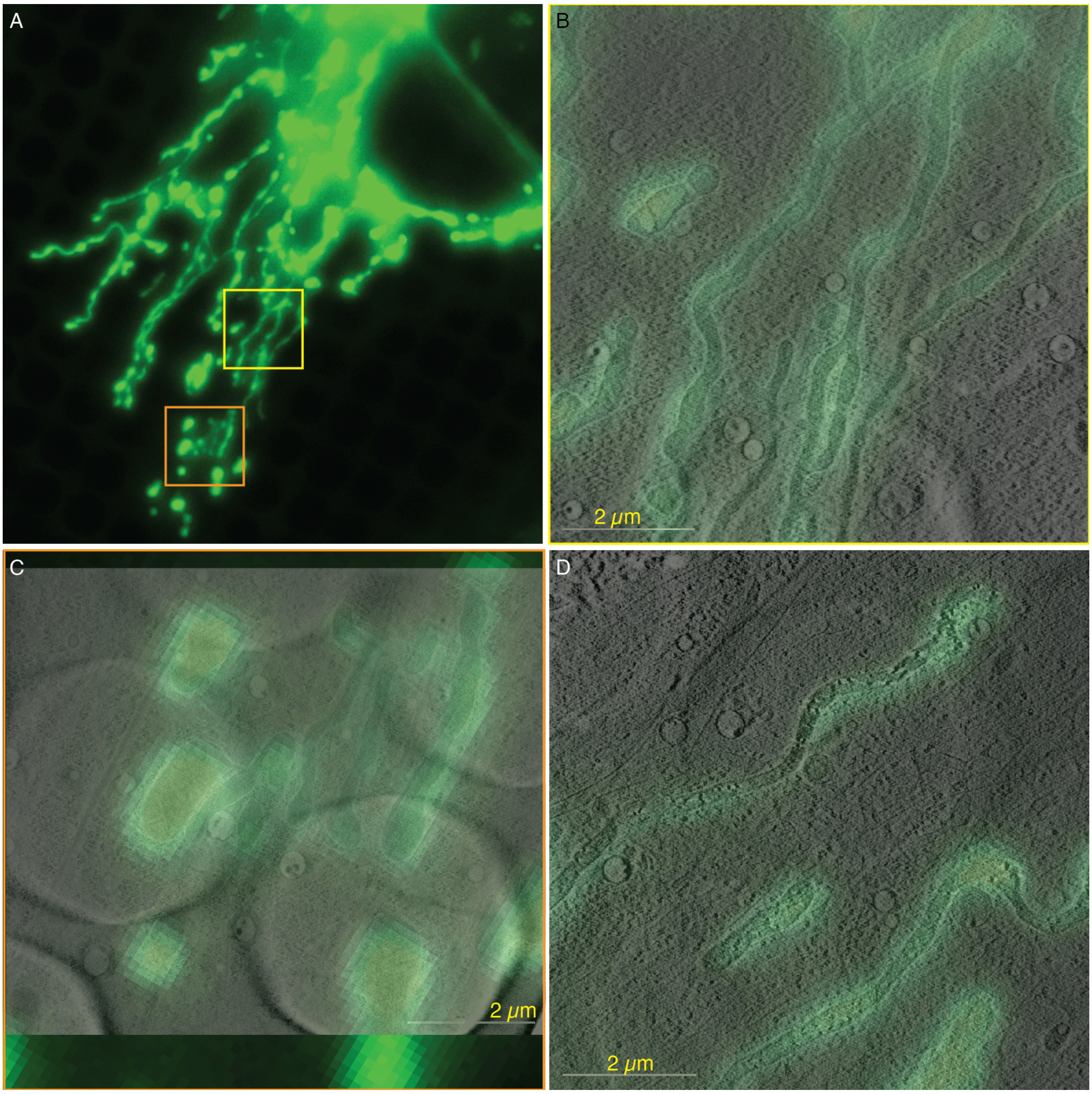


**Fig. S2.** CSTET data overlay on cryo-FM of unstressed MFF$^{-}$ mitochondria. The yellow and orange Box shows the area of the respective tomogram shown in (B) and (C) shown in Fig. S1. (D) was taken in a different cell on the same sample. The same slice is shown in Fig. 2A.

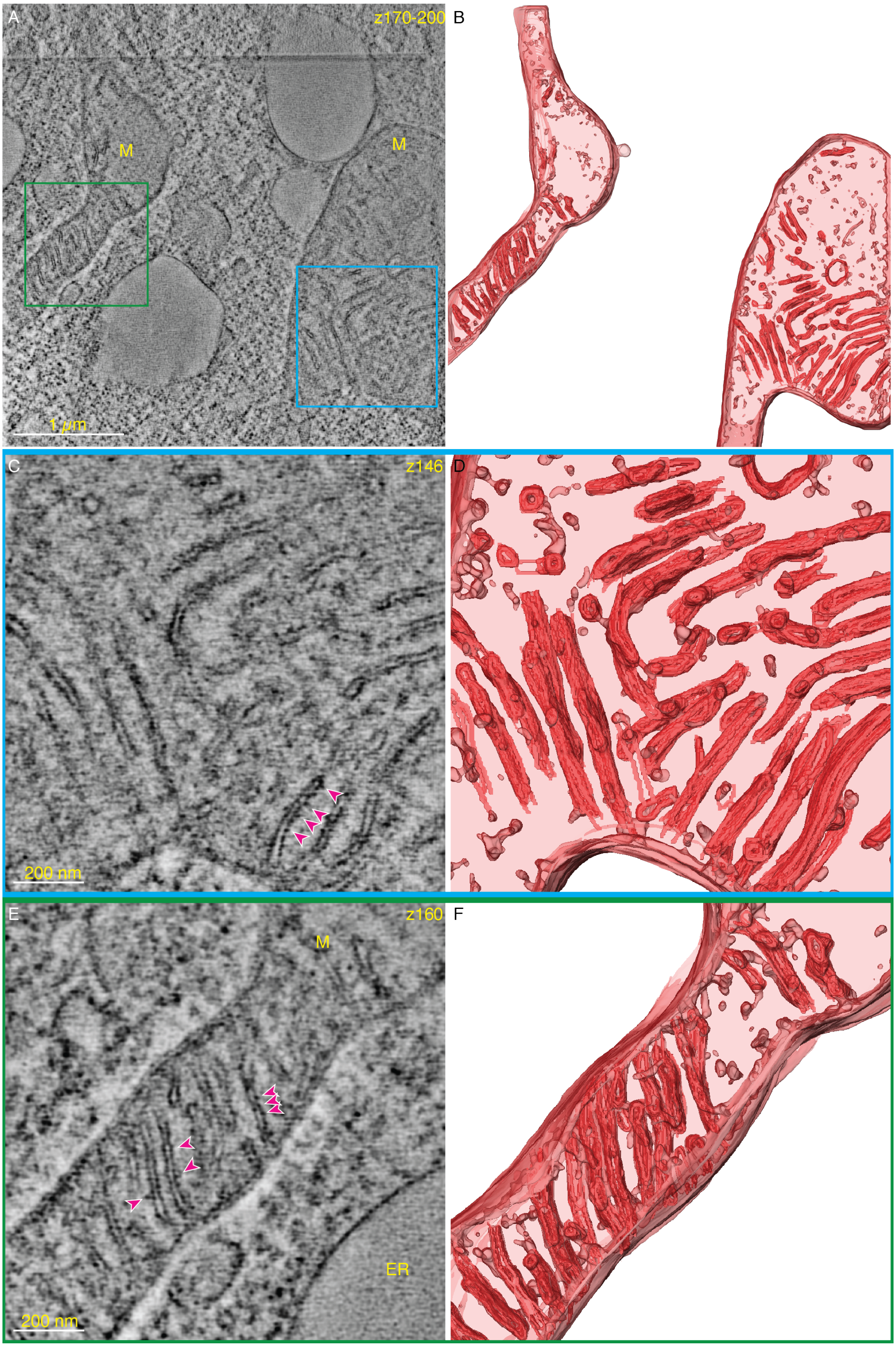


**Fig. S3.** A close up on the cristae network of unstressed MFF$^{-}$ mitochondria from Fig. 2B. (A) A 60 nm thick slice through the tomogram. Blue and green boxes show the area zoomed-in in (C+D) and (E+F), respectively. (B) 3D segmentation of the mitochondria in this tomogram. (C) Zoom-in into the cristae network (blue box). Scale bar is 200 nm. (D) 3D segmentation of the region. (E) Zoom-in into the cristae network (green box). Scale bar is 200 nm. (F) 3D segmentation of the region. Magenta arrowheads show protein densities in the cristae network.

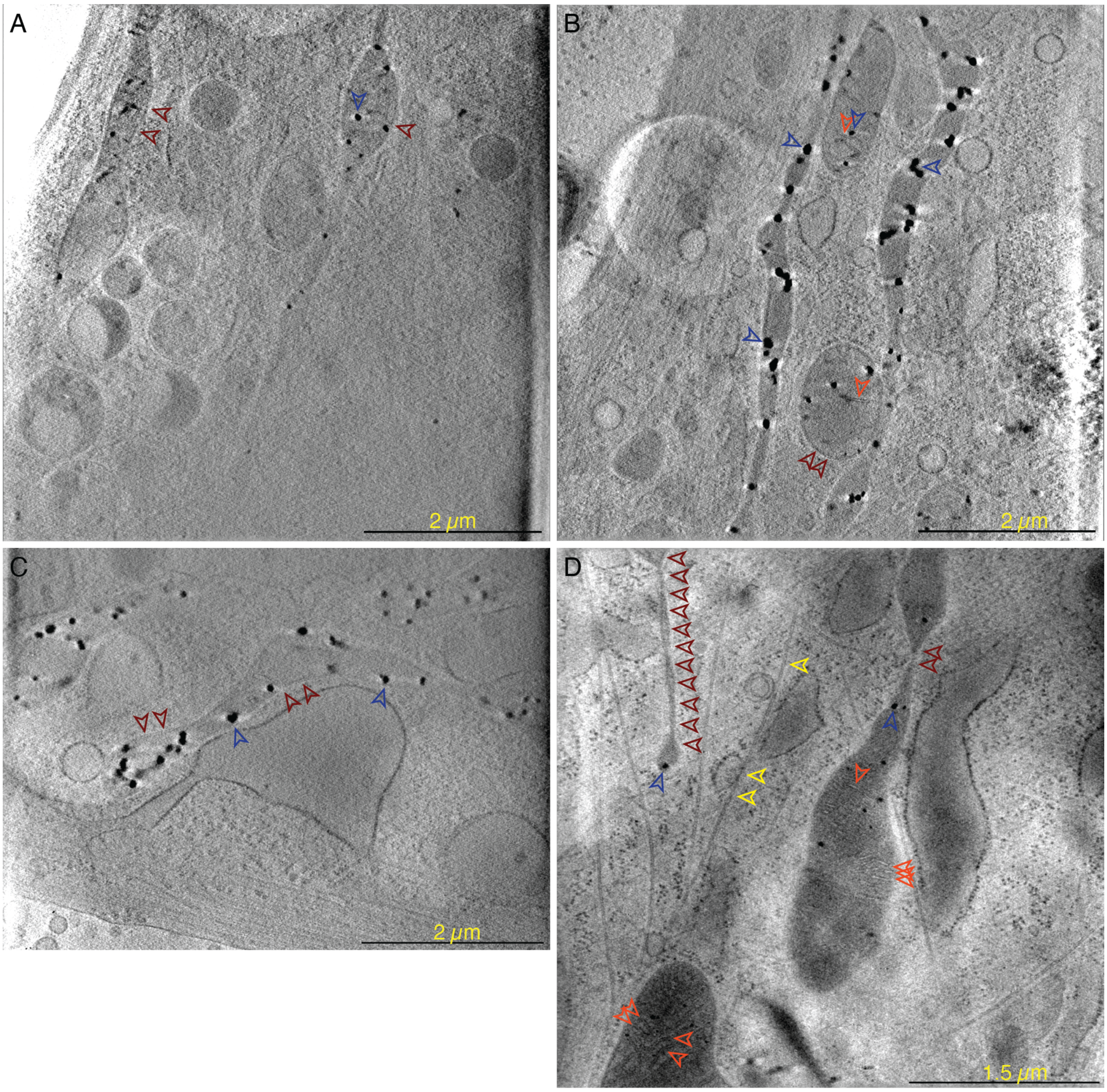


**Fig. S4.** MFF- cells under fission inducing conditions: A gallery of different constriction events. The calcium phosphate granules (dark blue), microtubules (yellow), cristae (light red) and OMM (dark red)are highlighted with arrowheads. Please note, these tomogram have not been deconvolved but have been filtered with the SIRTlike 30 filter in IMOD. Scale bars are 2 µm in (A-C) and 1.5 µm in (D). The circular structures, seen most clearly in (B), with a diameter of 2 µm, are holes in the carbon layer on the EM grid.

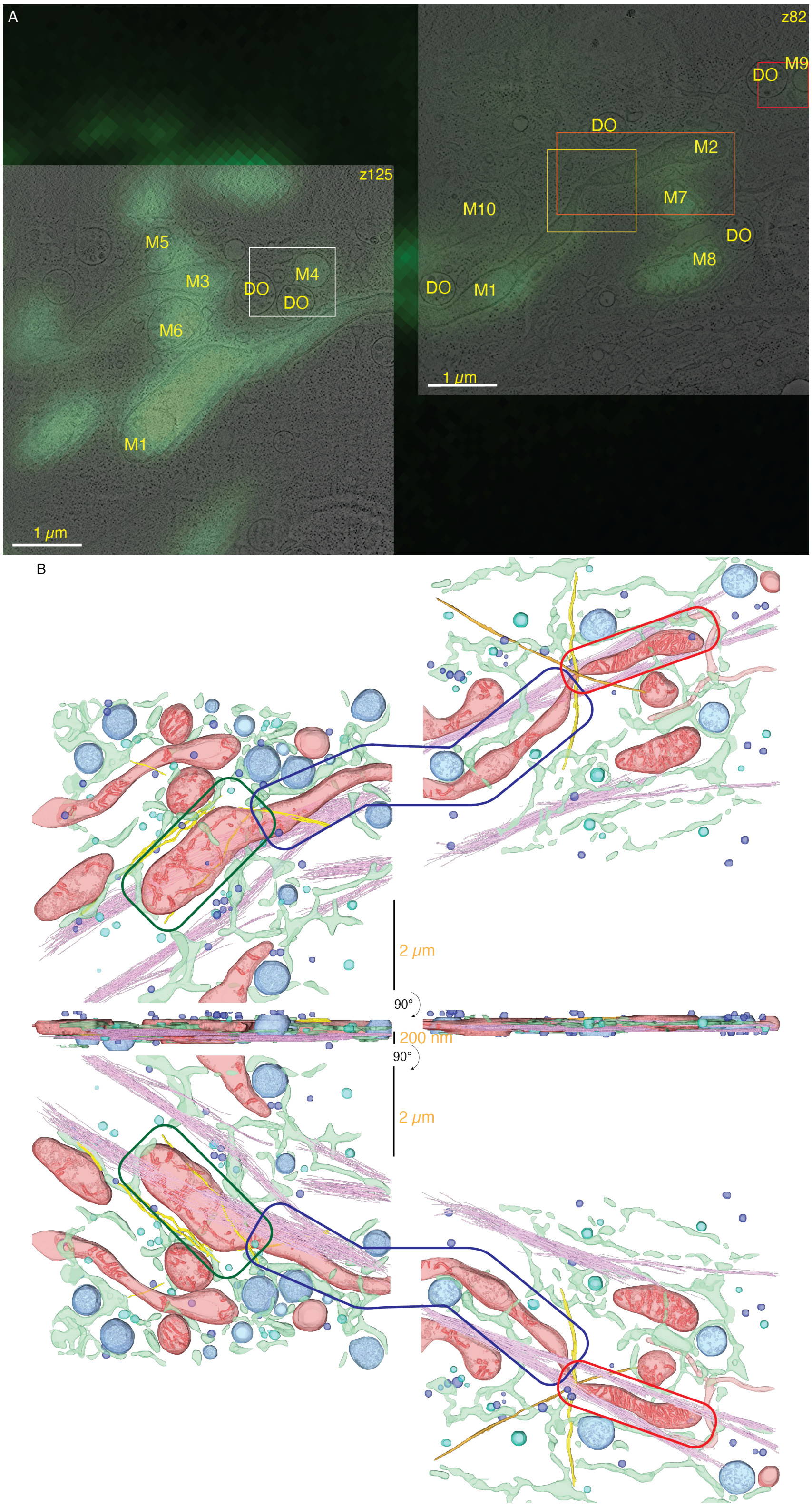


**Fig. S5.** Showing the spatial context to Fig. 6. (A) The tomograms from Fig. 6 are overlayed on the GFP channel of the cryo-FM image. The yellow and orange boxes show the area detailed in Fig. 7A and B, respectively. The red and white boxes show small mitochondrial fragments in contact with degradative organelles (Fig. 8 and **??**), but these are still green fluorescent. (B) Top, side, and bottom views of the 3D segmentation from Fig. 6. Green, blue and red boxes marks part one, two and three of the mitochondria, respectively. Red, green, yellow, orange, old pink, light red tube, dark blue and light blue mark the mitochondria, ER, microtubules, microtubules, actin, unknown tubular structure, microvesicles and degradative organelles.

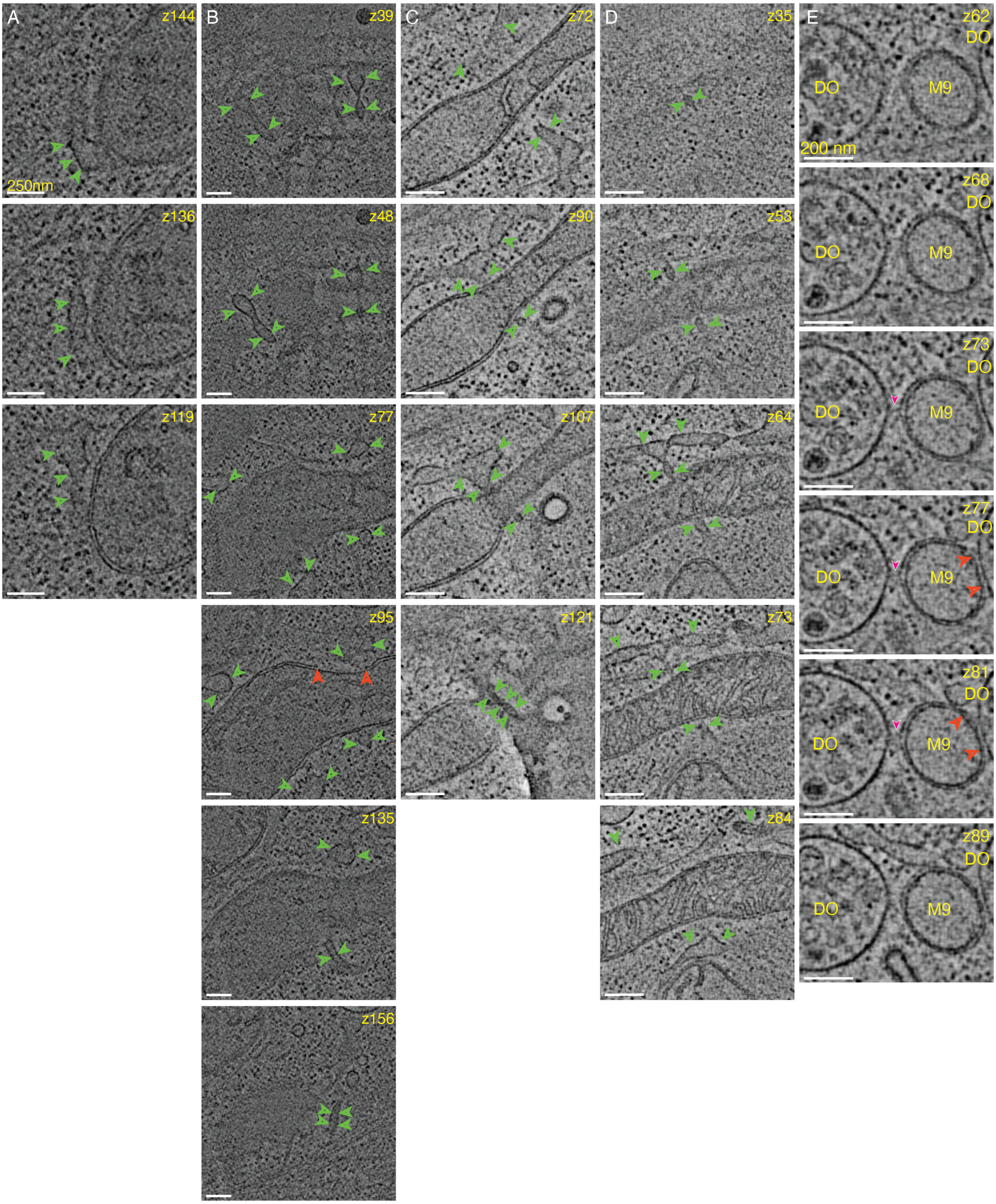


**Fig. S6.** Some detailed views of ER-mitochondria interactions from Fig. 6, and Fig. S5). Please note the arrangement follows along the mitochondria M1 from the tip towards M2. (A) shows areas near the tip of M1, (B) a part close to the edge between parts one and two of M1 (green and blue boxes in Fig. S5B). (C) Another ER below M1 on the edge of the carbon. (D) An ER tubule above M2. (E) A small mitochondrial fragment (M9) interacting with the digestive organelle (DO). See the red box in Fig. S5 for the area and the green fluorescence of the mitochondria. Red, magenta, and green arrowheads show the missing OMM, contact between M and DO, and ER, respectively. Scale bar is 250 nm in (A-D) and 200 nm in (E).

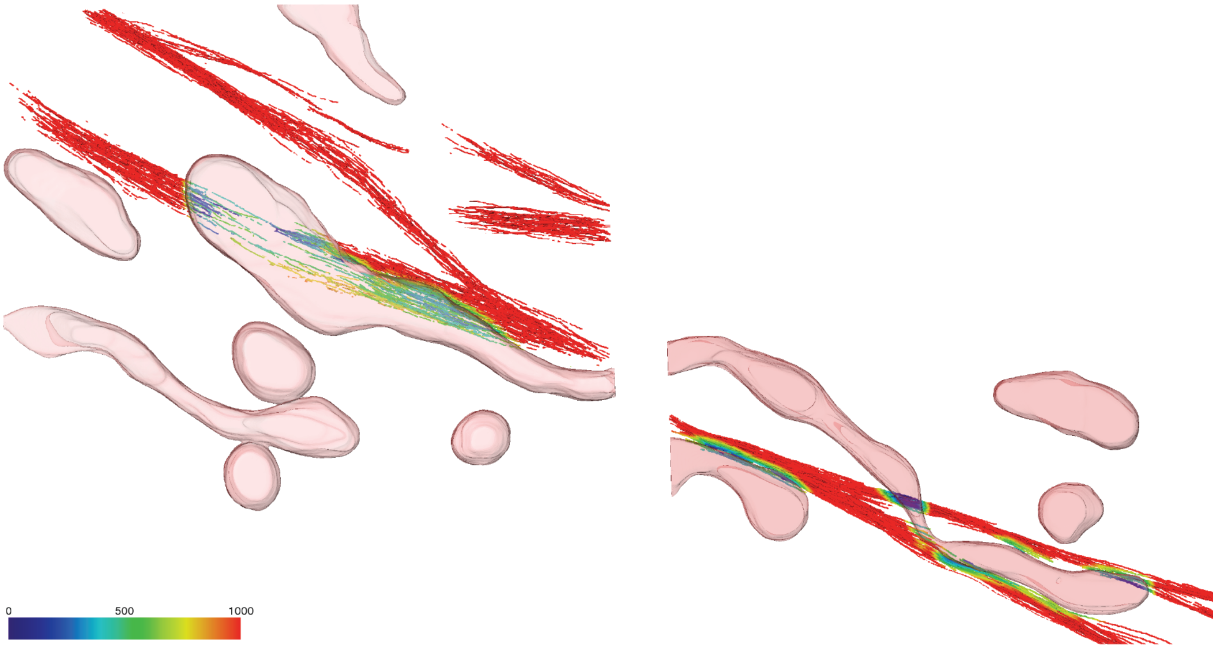


**Fig. S7.** Representation of the distance between the two actin bundles to the third part of the mitochondrion. The distance is represented in blue to red from 0 to >1000 nm.

#### 2 Observation summary

**Table S1.** Overview of the observations in MFF$^{-}$ cells under normal conditions

| | | | |
|---|---|---|---|
| Morphology | Elongated mitochondria with variable mitochondrial diameter | Hardly any tight constrictions | Mitochondrial derived vesicle formation |
| Cristae Pattern | Ordered cristae pattern | Small areas devoid of cristae | |
| Cellular objects in proximity | (ER-)Vesicles | | |
| Physiological observations | Calcium phosphate deposits can be missing | | |

**Table S2.** Overview of the observations in MFF$^{-}$ cells under fission inducing conditions

| | | | | |
|---|---|---|---|---|
| Mitochondrial morphology | "Beads-on-the-string" | Balloon- or bulklike mitochondria | Elongated mitochondria with variable diameter | |
| Cristae Pattern | No cristae | Degraded cristae | Ordered cristae pattern only in small portions | |
| Cellular objects in proximity | ER | Microtubules | Actin bundles | Degradative organelles |
| Physiological observations | Calcium phosphate deposits can be missing | | | |

#### 3 SerialEM dual-axis script

The script can be downloaded from Nexperion using this link: `https://serialemscripts.nexperion.net/script/81`

##### 3.1 How to run the script:

1. Before running the script save the navigator as a new file
2. Remove the objective aperture and return to the smallest magnification in LowMag-STEM mode
3. Change the TempFile name and location in the script
4. Open a new .mrc file (different from the TempFile)
5. Start the script and watch it run:
   5.1 The stage moves to the coordinates defined in the script (Variables: RotationcenterX and RotationcenterY)
   5.2 It takes a Record image, saves it in the open file and makes it a new map in the Navigator

5.3 It rotates the grid inside the microscope by 90° (set by the RotationTarget Variable in the script) and takes a new Record image at the coordinates defined in the script

5.4 The script calls the Align-with-rotation procedure with a search settings of 90°and plus-minus 6° (can be adjusted in the script)

5.5 All the items in the Navigator are then rotated by the amount calculated

5.6 The Record images are then saved in a new file

6. The new Navigator is now rotated at around 90° and when the position of the tomogram is not too close to the edge of the grid square automated continuation of the data collection including the 'realign-to-items' procedure works very well.
